# Description of *Schaedlerella arabinophila gen. nov., sp. nov.*, a D-arabinose utilizing bacterium isolated from feces of C57BL/6J mice and a close relative of *Clostridium* sp. ASF 502

**DOI:** 10.1101/496661

**Authors:** Melissa Soh, Sou Miyake, Austin Lim, Henning Seedorf

## Abstract

The use of gnotobiotics has gained large interest in recent years due to technological advances that have revealed the importance of host-associated microbiomes for host physiology and health. One of the oldest and most important gnotobiotics mouse model, the Altered Schaedler Flora (ASF) has been used for several decades. ASF comprises eight different bacterial species, which have been characterized to different extent, but only few are available through public strain collections. Here, the isolation of a close relative to one of the less studied ASF strains, *Clostridium* sp. ASF 502, is reported. Isolate TLL-A1, which shares 99.6% 16S rRNA gene sequence identity with *Clostridium* sp. ASF 502, was obtained from feces of C57BL/6J mice where is was detectable at a relative abundance of less than one percent. D-arabinose was used as sole carbon source in the anaerobic cultivation medium. Growth experiments with TLL-A1 on different carbon sources and analysis of its ~6.5 gigabase genome indicate that TLL-A1 harbors a large gene repertoire to utilize different carbohydrates for growth. Comparative genome analyses of TLL-A1 and *Clostridium* sp. ASF 502 reveal differences in genome content between the two strains, in particular with regards to carbohydrate activating enzymes. Based on physiology and genomic analysis it is proposed to name TLL-A1 to *gen. nov. sp. nov Schaedlerella arabinophila* TLL-A1 (DSMZ 106076^T^; KCTC 15657^T^). The closely related *Clostridium* sp. ASF 502 is proposed to be renamed to *Schaedlerella arabinophila* to reflect its taxonomic standing and to keep ‘ASF 502’ as strain designation.

**Importance:** The Altered Schaedler Flora (ASF) remains an important tool to mechanistically investigate host-microbe interactions in the mammalian alimentary tract. Extensively characterizing the eight different bacterial strains, which are constituting ASF, has the potential to further increase the definition of this widely used model microbial community and to enhance our understanding of how individual microorganism interact with a host and/or how they may affect its physiology. However, some of the ASF strains have unfortunately been lost or are not easily accessible to the scientific community. The isolation and characterization of the here described species, proposed to be named *Schaedlerella arabinophila*, which is closely related to *Clostridium* species ASF502, may therefore be an important corner stone in further improving the value of studies using ASF or other defined synthetic microbial communities that require usage of autochthonous microorganisms in gnotobiotic mice.

## Introduction

Microbiome studies have revealed extensive insights into the complex associations of microorganisms with specific phenotypes of host physiology and health (1, 2). A major focus of the studies remains on the mammalian intestinal tract, which is densely populated with microorganisms from all three domains of life (3, 4). However, it is often not well established whether an association of microbial taxa (or a collection thereof) with a host phenotype could also equate to being the underlying cause. Establishing this causality still often relies on the use of model organisms, such as the laboratory mouse, which offers several advantages: they are well-characterized, can be genetically modified, and can be reared germ-free. The latter is of particular interest as it allows to perform experiments with gnotobiotic animals (germ-free animals that are associated with defined microbiota). The use of gnotobiotic mice was already recognized in the middle of the last century when it became possible to rear germ-free mice and to associate them with autochthonous microorganisms (5). One of the most prominent examples from this era is the Altered Schaedler Flora (ASF). ASF is a defined microbial community consisting of eight bacterial species, and has been used for a large number of studies that investigate host-microbe interactions (6–8). ASF is an updated form of the ‘Schaedler flora’ and was revised by Orcutt *et al*. in 1978 to standardize its use in gnotobiotic experiments (9, 10). At the time, the strain characterization and selection was mostly based on physiology and microscopy from the cultivatable fraction of the enteric microbial community, given that sequence-based characterization of microorganisms was still in its infancy. Subsequent phylogenetic analyses of the 16S rRNA genes enabled additional taxonomic identification of the strains, and thus the strains were named ASF 356 (*Clostridium* species), ASF 360 (*Lactobacillus intestinalis*), ASF 361 (*Lactobacillus murinus*), ASF 457 (*Mucispirillum schaedleri*), ASF 492 (*Eubacterium plexicaudatum*), ASF 500 (*Pseudoflavonifractor* species), ASF 502 (*Clostridium* species) and ASF 519 (*Parabacteroides goldsteinii*). See Dewhirst *et al*. for more details on phylogenetic information of each ASF strain (11). Meanwhile, all eight strains of ASF were genome sequenced (to draft quality) which have further characterized them and have added value to using ASF as model community in gnotobiotics experiments (12).

However, the amount of information available on physiology and ecological role varies greatly between each of the ASF strains, as some of them remain difficult to cultivate. Detrimental to further characterization efforts is the fact that some of the original ASF strains are not available (anymore) through public strain collections (11), and are therefore not easily accessible. In some cases (e.g. ASF 492 (*Eubacterium plexicaudatum*), the original strain has been lost (11). Nevertheless, ASF remains a valuable model, and close relatives have been isolated for some of the lost strains. One recent example is the isolation and formal description of *Mucispirillum schaedleri* ATCC BAA-1009^T^, a close relative of ASF 457 (13). The type strain has undergone additional characterizations that provide insights into the lifestyle of this species in the mucus layer of the mouse intestinal tract (14), which may also have implications for interpreting the role of ASF 457 in ASF and the use of ASF in general. For other strains, such as ASF 502, less information is available. A qPCR-based study indicated that the abundance of this particular strain can be highest of all eight ASF strains in ASF-harbouring mice feces (15). The same study also revealed that the abundance level of the strain varied considerably in the different sections of the intestinal tract, highlighting that this species may have important roles in specific segments of the intestinal tract of ASF-harbouring mice (15).

Here, we present the isolation and formal description of a new genus and species, *Schaedlerella arabinophila* TLL-A1. *S. arabinophila* TLL-A1 was isolated from feces of C57BL/6J mouse and its 16S rRNA gene shares 99.6 percent sequence identity with that of *Clostridium* species ASF 502. Concurrent to phenotypic and genomic characterization, 16S rRNA-amplicon sequencing was performed to determine the relative abundance of *S. arabinophilus* in healthy mouse fecal samples. Lastly, the genome of *S. arabinophila* was sequenced and compared to the available draft genome of *Clostridium* species ASF 502 to identify differences between the two strains in genome content.

## Results

### Isolation and whole genome analysis of *S. arabinophila* TLL-A1

*S. arabinophila* TLL-A1 was isolated as part of a larger cultivation effort that screened for utilization of specific carbon sources by mouse intestinal microorganisms. For convenience, the isolated strain TLL-A1 and the closest known relative *Clostridium* sp. ASF 502 (GenBank ID GCA_000364245.1) will henceforth be referred to as TLL-A1 and ASF 502, respectively. The colonies of TLL-A1 were initially obtained on solid medium containing 1.6 g per litre D-arabinose. The observed colonies appeared to be uniform in shape and colour. Ten colonies were selected for growth in liquid medium and DNA was extracted from cultures. No other microbial species were detected on clone libraries and the obtained strains (all identical) had highest sequence identity to ASF 502. The purity of cultures was confirmed by serial plating and selection of colonies before inoculating into larger liquid medium to obtain sufficient cell mass for genome sequencing.

The gDNA was sequenced on PacBio RSII and raw reads were assembled by HGAP 4 with further error correction conducted by Pilon. The Pilon assembly yielded 3 contigs for a total assembly size of 6,521,014 bp, with G+C content of 47.9%, N50 of 6,346,997, and an average coverage of 107x across the genome. The large N50 is due to the three contigs having uneven size (6.35 Mb, 0.16 Mb and 9.5 kb). The two shorter contigs were not identified as plasmids with Plasmid Finder (16). Reads from MiSeq paired-end sequencing was used for further error correction using Pilon (17). Because the assembly by SMRT analysis was already of high quality, Pilon only corrected for 10 locations totalling 11 bases. The total assembled genome was 6.52 Mb, comparable to 6.36 Mb draft genome published for *Clostridium* sp. ASF 502 (12). Circular display illustrates the three contigs, CDS on forward and reverse reads, non-CDS features, AMR genes, transporters as well as GC content and GC skew (Figure 1). The genome has an estimated completeness of 94.7% and contamination of 2.44% based on 333 marker genes conserved in Lachnospiraceae as identified by checkM (Table S1). The assembly contain 6,724 CDSs; of which 59 were RNAS (6 rRNA and 53 tRNA genes), 313 repeat regions, 23 and 22 CRISPR repeat and spacer regions, respectively. There are 3,225 genes (48.0%) assigned putative functions and 3,499 hypothetical proteins. 3,632 are assigned as FIGfam. Table S2 summarises the genome statistics. All annotations are publically available online (www.patricbrc.org) under genome ID 97139.17.

**Figure 1.**
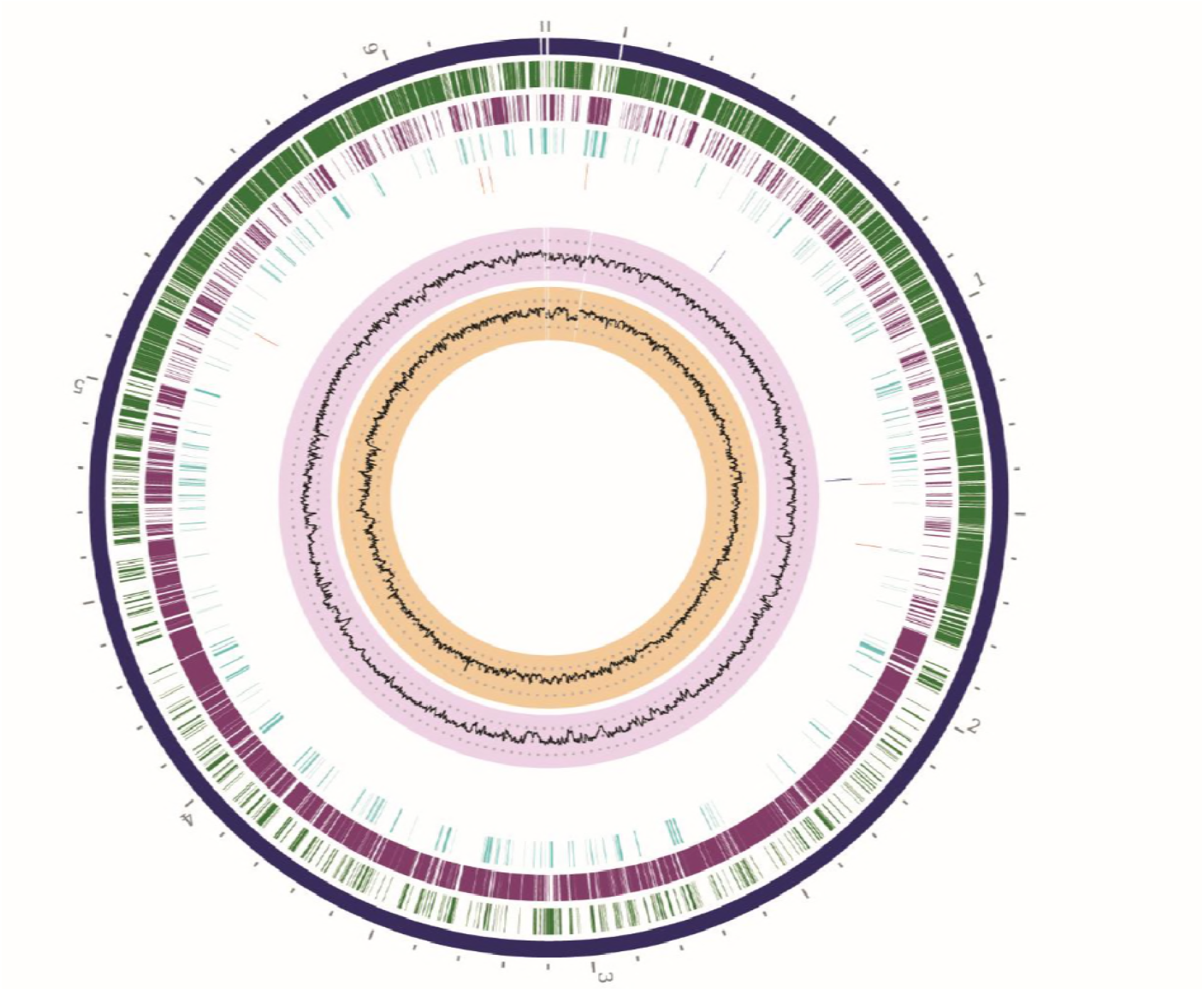
Circular display of *Schaedlerella arabinophila* strain TLL-A1 genome displaying (from outside to in) contiguity, CDS on forward and reverse reads, non-CDS features, AMR genes, transporters as well as GC content and GC skew.

### 16S rRNA gene and whole-genome phylogeny of *S. arabinophila*

16S rRNA gene phylogeny of *S. arabinophila* strain TLL-A1 and closely related *Clostridium* strain ASF 502 (GenBank ID GCA_000364245.1) show that two are closely related, with *Clostridium hylemonae* and *Clostridium scindens* the closest known relatives outside the clade (Figure 2A). This is in stark contrast to whole-genome phylogeny where TLL-A1 and ASF 502 are clustered with *C. glycyrrhizinilyticum* and *Ruminococcus* species (Figure 2B). Note that the difference may be due to bias in the availability of 16S rRNA gene and whole genomes. At 16S rRNA gene level, TLL-A1 and ASF 502 shared 99.6% sequence identity. Both average amino acid identity (AAI) and genome sequence-based delineation (GGDC) confirmed two strains to have high probably of belonging to the same species (Two-way AAI: 94.46% (SD: 14.50%) from 3976 proteins; Probability that DDH > 70% = 92%, respectively). Given that the two strains are considered the same species, we then opted to determine their phylogenetic relationship with the closest sequenced relative, *Clostridium glycyrrhizinilyticum* JCM 13369 (NCBI accession no. NZ_BBAB00000000). GGDC showed that the probability that *C. glycyrrhizinilyticum* and TLL-A1 belonged to the same species (i.e. DDH > 70%) was 0% (via logistic regression). At 16S rRNA gene level, the two species shared 93.25% sequence identity indicating that TLL-A1 and ASF 502 together likely represent a new genus. Thus, strains TLL-A1 and ASF 502 were named *Schaedlerella arabinophila* TLL-A1 and *Schaedlerella arabinophila* ASF 502, respectively (formal description below).

**Figure 2.**
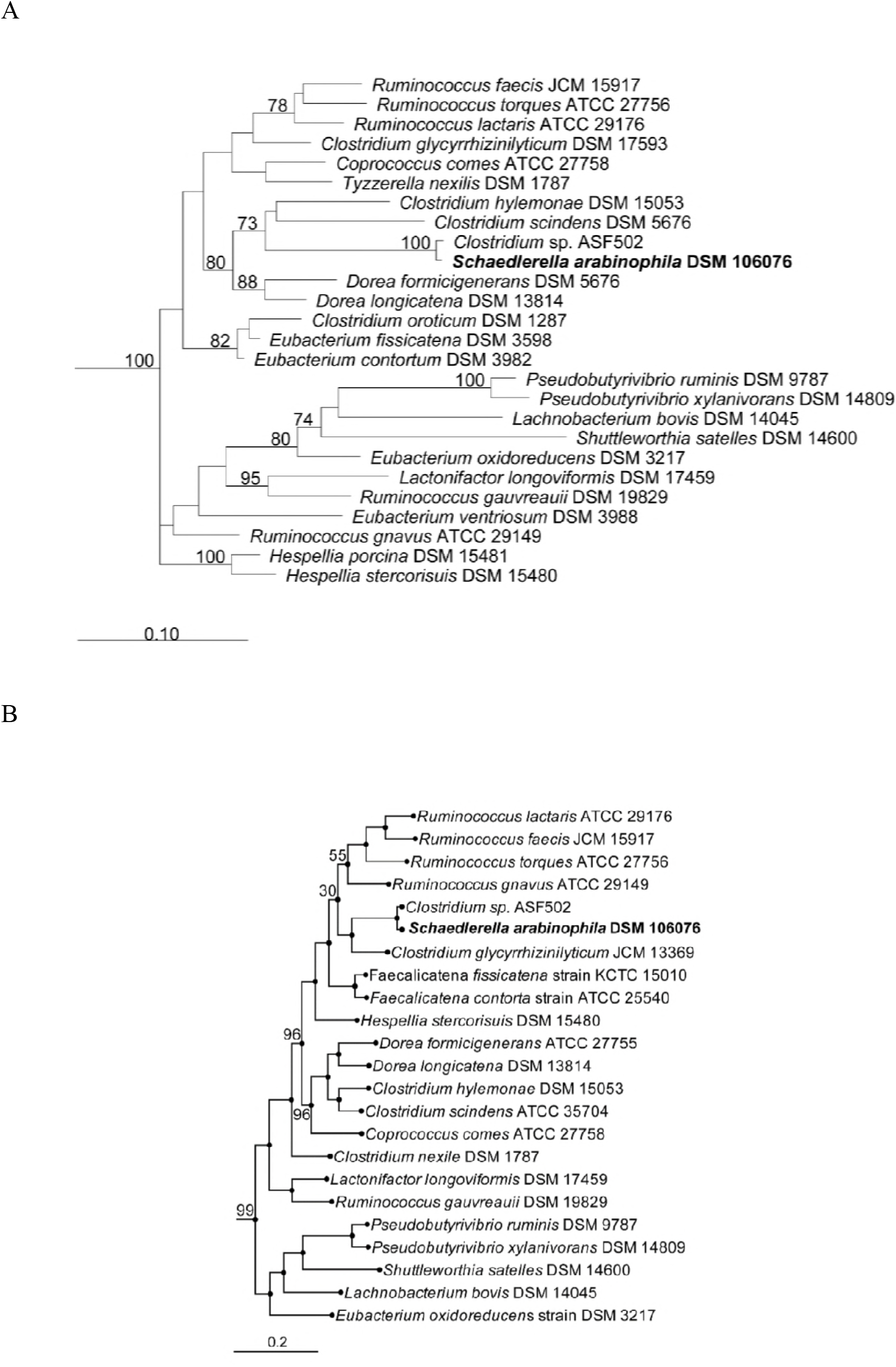
Phylogenetic trees showing the relationship of *Schaedlerella arabinophila* to closest related bacteria. A) RaxML 16S rRNA gene phylogeny of *Schaedlerella arabinophila* strain TLL-A1 (in bold) and related organisms. The sequences were SINA-aligned (Pruesse et al., 2012) and the tree was constructed in ARB. B) Whole-genome phylogeny of *Schaedlerella arabinophila* and related genomes. The tree was constructed in PATRIC with RaxML algorithm.

### Comparative genome analysis of *S. arabinophila* with *Clostridium* sp. ASF 502

Genome comparison indicated similar general parameters for the two strains (Table S2). In addition, overall differences in the genomes were compared using clusters of orthologous genes (COGs). Of 4,226 and 4,257 COGs identified in TLL-A1 and ASF 502, 4081 were shared between the two, which meant that 145 (3.43%) and 176 (4.13%) were unique to TLL-A1 and ASF 502, respectively (Table S3 and Table S4). For the unique clusters, whist most of the Swiss-Prot annotation remained unclassified, those given Gene Ontology (GO) terms were grouped according to their molecular functions (Figure 3). TLL-A1 had higher occurrence of unique genes related to cofactor binding (GO:0048037), electron carrier activity (GO:0009055), hydrolase activity (GO:0016787), ion binding (GO:0043167), isomerase activity (GO:0016853), metal cluster binding (GO:0051540), nucleotide binding (GO:0000166), nutrient reservoir activity (GO:0045735), oxidoreductase activity (GO:0016491), transferase activity (GO:0016740) and transporter activity (GO:0005215); while ASF 502 had more of catalytic activity (GO:0003824), lyase activity (GO:0016829), molecular function (GO:0003674), protein binding (GO:0005515) and signal transducer activity (GO:0004871) genes. In particular, electron carrier activity (GO:0009055), metal cluster binding (GO:0051540) and nutrient reservoir activity (GO:0045735) were exclusive to TLL-A1 while catalytic activity (GO:0003824), lyase activity (GO:0016829) and protein binding (GO:0005515) were only found in ASF 502 (Figure 3).

**Figure 3.**
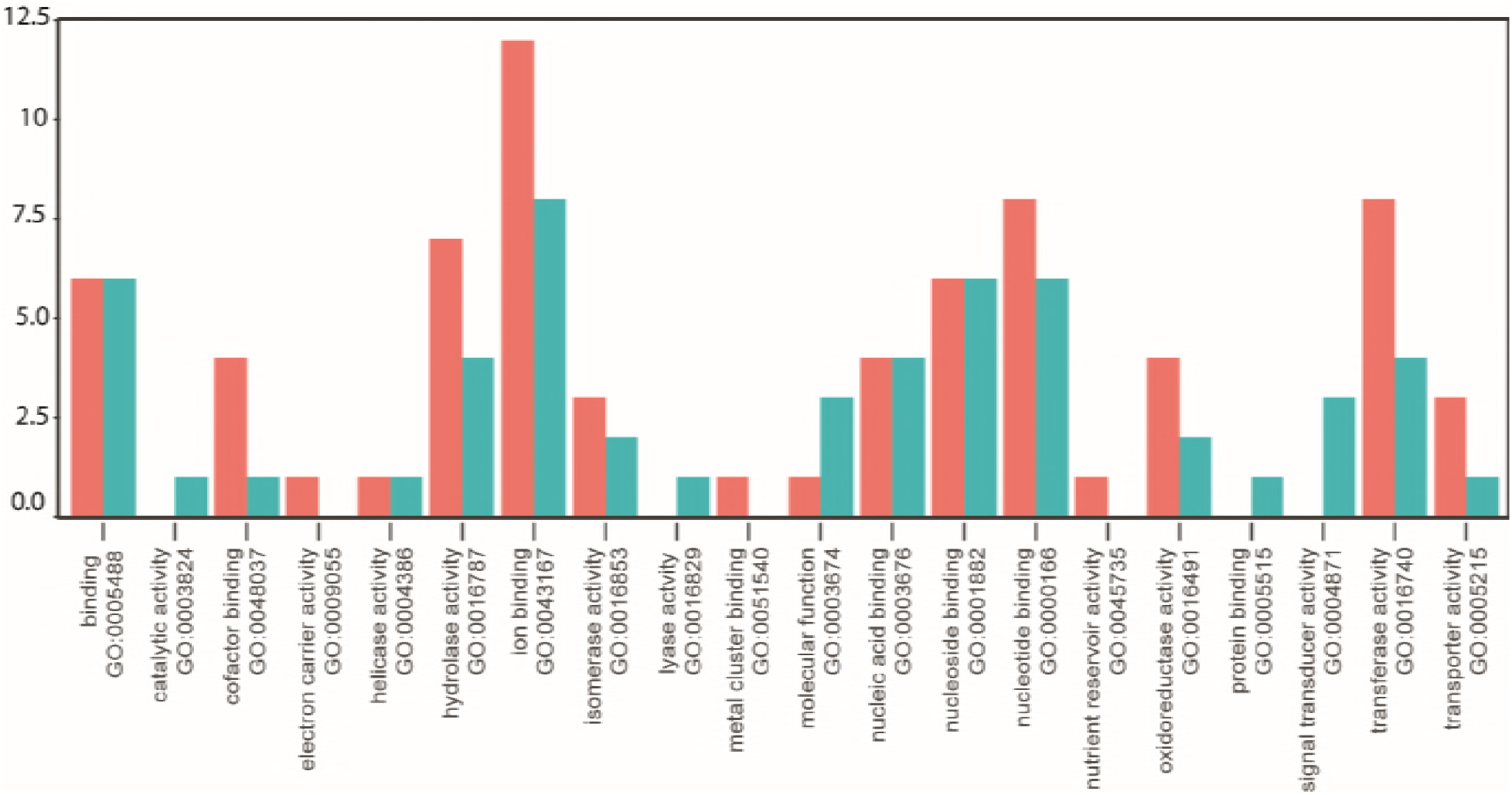
Differences in predicted molecular function between *Schaedlerella arabinophila* strains TLL-A1 and ASF 502. Shown are counts of Swiss-Prot annotations of unique COGs in *Schaedlerella arabinophila* TLL A1 (red) and ASF 502 (green) classified into Gene Ontology (GO) terms according to molecular function.

Carbohydrate Active EnZymes (CAZymes) are set of enzymes involved in the synthesis, breakdown and transport of the carbohydrates. Given the difference observed in the unique portions of the COGs, we further compared the CAZymes between the two genomes to see if there are any clear differences in the carbohydrate-degrading enzymes. The strain TLL-A1 genome encodes 168 CAZymes, which were classified as 124 glycoside hydrolases (GHs), 30 glycosyl transferases (GTs), 7 carbohydrate esterases (CEs) and 7 carbohydrate-binding modules (CBMs), while ASF 502 genome consisted of 130 CAZymes, classified into 90 GHs, 28 GTs, 6 CEs and 6 CMBs (Figure 4). Family level classification of GHs showed both genomes comprised of similar enzymes, except TLL-A1 had higher counts of GHs. β-glucosidase/ β-galactosidase, responsible for cleaving oligosaccharides, were the most abundant enzymes found in both genomes, comprising 7 families (GH1, 2, 3, 31, 35, 36, 42) in varying number of genes. Other abundant GHs included GH78 (α-L-rhamnosidase (EC 3.2.1.40), 13 and 6 in TLL-A1 and ASF 502 respectively), GH13 (α-amylase (EC 3.2.1.1), 10 and 7), GH39 (α-L-iduronidase (EC 3.2.1.76), 9 and 5) and GH43 (β-xylosidase (EC 3.2.1.37), 8 and 5). Interestingly, there were only three unique GHs in either of the genomes, which were GH130 (β-1,4-mannosylglucose phosphorylase (EC 2.4.1.281)) and GH 136 (lacto-N-biosidase (EC 3.2.1.140)) found in TLL-A1 and GH20 (β-hexosaminidase (EC 3.2.1.52)) found in ASF 502.

**Figure 4.**
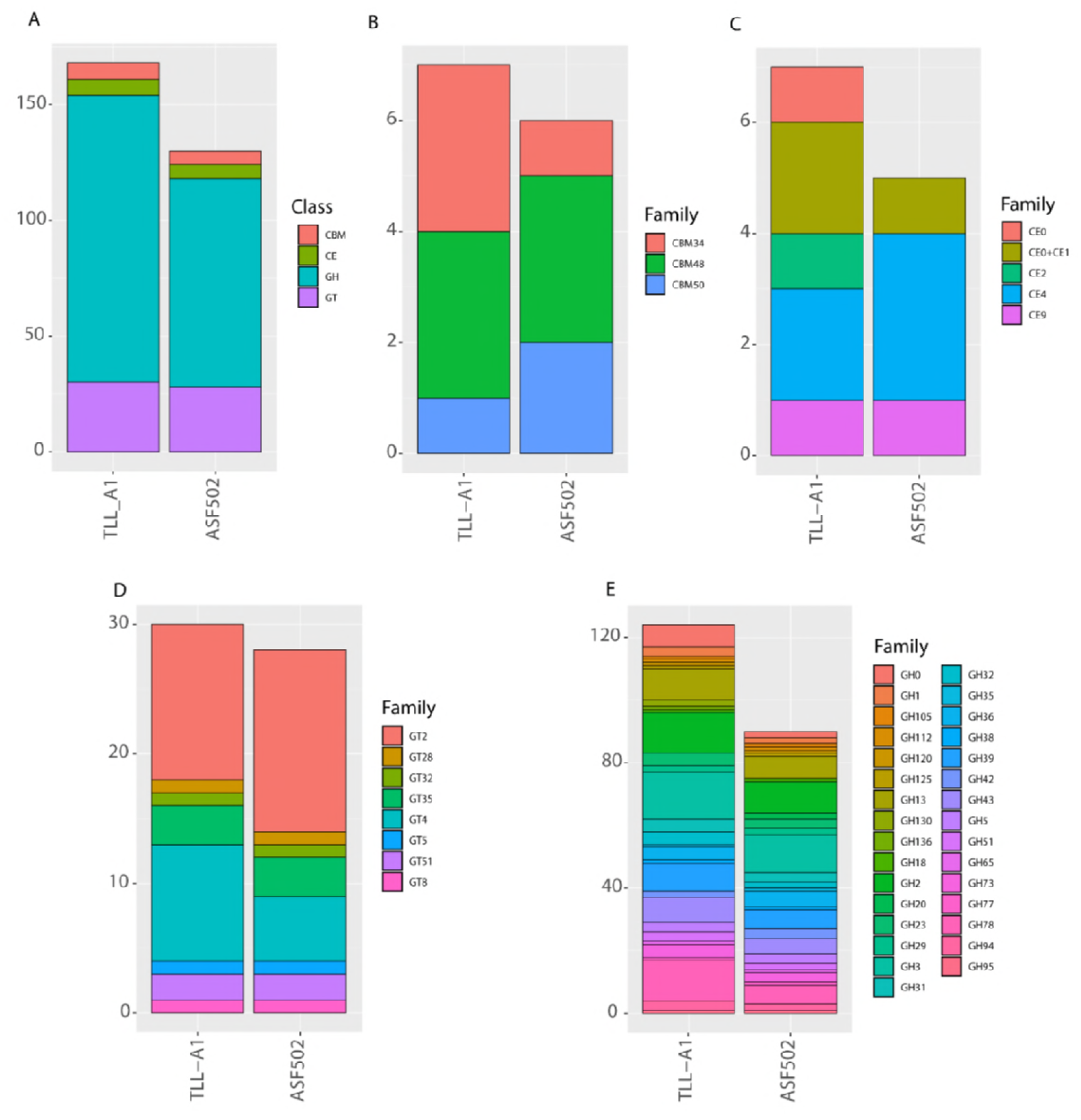
Comparative analysis of carbohydrate Active EnZymes (CAZymes) identifies differences between *Schaedlerella arabinophila* strain TLL-A1 and ASF 502 in the predicted capability to process carbohydrates. A) Overall comparison of CAZyme category counts between strain *S. arabinophila* strain TLL-A1 and ASF 502. B) Family level classification of CBMs in *Schaedlerella arabinophila* and ASF 502. C) Family level classification of CEs in *Schaedlerella arabinophila* and ASF 502. D) Family level classification of GTs in *Schaedlerella arabinophila* and ASF 502. E) Family level classification of GHs in *Schaedlerella arabinophila* and ASF 502. Only CAZymes identified with > 2 methods (out of HMMER (E-Value < 1e-15, coverage > 0.35; http://hmmer.org/), DIAMOND (E-Value < 1e-102; (47)) and Hotpep (Frequency > 2.6, Hits > 6; (48)) were considered, as recommended by the authors.

Distribution of GTs were also similar between TLL-A1 and ASF 502 genomes. GT2 (cellulose synthase (EC 2.4.1.12) / chitin synthase (EC 2.4.1.16)) was the most abundant, followed by GT4 (sucrose synthase (EC 2.4.1.13)), GT35 (glycogen or starch phosphorylase (EC 2.4.1.1)) and GT51 (murein polymerase (EC 2.4.1.129)). In comparison to GHs and GTs, CEs and CBMs were both less abundant, with only three families identified each (CE2, 4, 9; CBM34, 48, 50).

### Physiological properties of *S. arabinophila strain* TLL-A1

In terms of physical characteristics, TLL-A1 was found to be Gram-negative and rod shaped. Doubling time under experimental condition was ~6.5h and cells reached a length of three to five μm at the end of exponential/early stationary phase. Light microscopy did indicate refractive structures at the terminal section in a smaller subset of observed cells. However, spore staining was also not conclusive as only few cells seem to harbor potential spores.

The isolation medium for TLL-A1 contained D-arabinose as main carbon source, but additional carbon sources (listed in the methods section) were tested as growth substrates for this strain (see Table S5 for results). Overall, the growth experiments suggest that *S. arabinophila* strain TLL-A1 can utilize a wide variety of different carbon sources.

Results from the characterizations performed at DSMZ are given below and in the corresponding tables. Analysis of polar lipids indicates primary presence of phospholipids, glycolipids and phosphoglycolipids (Figure S1). Analysis of cellular fatty acids indicated high amounts of saturated C14 and C16 fatty acids (Table S6). API Rapid ID 32A revealed enzyme activity for most tested carbohydrates, but not for any of the tested amino acids (Table S7). No respiratory quinones were detected. Additionally, analysis of peptidoglycan total hydrolysate contained the following amino acids: meso-diaminopimelic acid, alanine and glutamic acid (additional proteinogenic amino acids were disregarded as protein contamination). In addition to the three abovementioned amino acids, the peptidoglycan partial hydrolysate contained the peptides L-Ala-D-Glu and Dpm-D-Ala. The peptidoglycan type of TLL-A1 was concluded to be A1y meso-Dpm-direct.

### Abundance of *S. arabinophila* TLL-Al in the feces of mice

DNA extracted from fecal pellets of four healthy male mice (strain C57BL/6J) were subjected to amplicon sequencing to determine the relative abundance of TLL-A1 in the original source of isolation. The gut microbiota of four 24-week old mice strain C57BL/6J were variable yet consisted mainly of recently re-classified *Muribaculum* (formerly known as S24-7 group), averaging 36.16% ±0.042 across four mice gut (Figure 5). Other abundant genera included *Faecalibaculum* (11.44% ±0.078), *Lactobacillus* (8.95% ± 0.062), *Turicibacter* (8.59% ±0.024) and a couple of Lachnospiraceae genera (6.28% ±0.015 and 4.12% ±0.016). *S. arabinophila* belonged to the Lachnospiraceae UCG-006 clade, which ranged in abundance from 0.87 – 0.11%. In order to know the relative abundance of TLL-A1, all sequences belonging to UCG006 was picked out and re-classified with custom database that included only UCG006 and TLL-A1 sequences. Sequences belonging to TLL-A1 were present in all four mice, yet they were not very abundant; ranging from 6 - 23% of all UCG-006 picked out.

**Figure 5.**
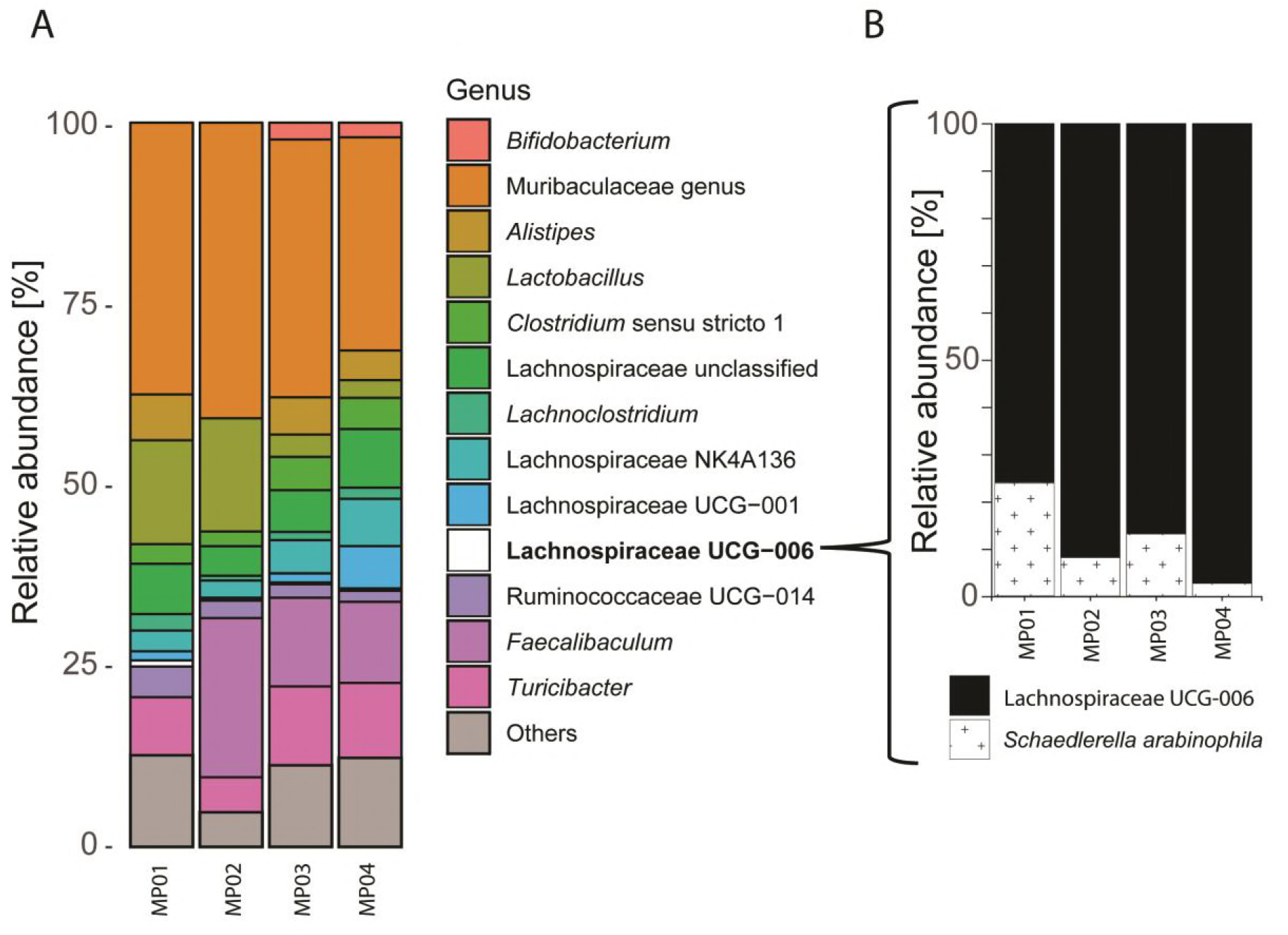
Analysis of relative abundance of *S. arabinophila* in the fecal microbiota of conventionally-raised C57BL/6J mice. A) Genus-level taxonomy composition of four healthy mice fecal microbiota (strain C57BL/6J). Genera that have a relative abundance of less than 1% within the sample are classified as others. B) Relative abundance of *Schaedlerella arabinophila* within Lachnospiraceae UCG-006, closest match in terms of 16S rRNA gene from Silva database. Fecal samples from same mice were used as inoculum for isolation of *S. arabinophila*

## Discussion

The initial aim of the study was to identify murine intestinal bacteria that have the capacity to utilize D-arabinose for growth. The cultivation experiments led to the isolation of a microbial isolate, named *Schaedlerella arabinophila* TLL-A1, which shared highest sequence identity with *Clostridium* species ASF 502, renamed here *Schaedlerella arabinophila* ASF 502. The isolation approach and characteristics of *S. arabinophila*, genomic differences between the two *S. arabinophila* strains, as well as the abundance of *S. arabinophila* in the mouse intestinal tract are discussed in the following.

### Isolation and characteristics of *Schaedlerella arabinophila* strain TLL-A1

Intestinal microorganisms are known to be efficient utilizers of a wide variety of different carbohydrates (18–20). The pentose arabinose is in this regard of particular interest as L-arabinose is more common in nature than D-arabinose, opposed to most other monosaccharides where the D-configuration is more common than the L-configuration. The use of D-arabinose as sole carbon source in solid medium resulted in growth of TLL-A1 in two independent experiments, with no other microbial species detected from the resulting colonies. Although the total number of selected colonies may be too small to be conclusive about how selective the medium is towards TLL-A1 isolation, the results indicate that either the growth conditions are only suitable for TLL-A1, or that the diversity of D-arabinose utilizers in the murine gut is low. The exact mechanism of D-arabinose utilization by TLL-A1 remains to be elucidated through follow-up analysis, but it is has been shown for other organisms, such as *E. coli*, that growth may occur via a L-fucose degradation pathway (21). Considering that fucose is a common component of host glycans (22), together with the finding that the *S. arabinophila* genomes encode for a large repertoire of CAZymes, it seems plausible that *S. arabinophila* may be able to utilize a similar pathway for growth. Additional characterization of the growth properties of *S. arabinophila* revealed that this species is able to utilize not only D-arabinose, but also a wide variety of other carbohydrates, which may indicate that this species is a versatile commensal in the murine intestine that may be able thrive on carbohydrates from different dietary sources.

The TLL-A1 genome also revealed the presence of sporulation genes (48 annotated genes related to sporulation). TLL-A1 cultures were therefore inspected for spores, but the results were inconclusive: initial light microscopy and spore staining revealed a few potential spores, but only in small numbers (less than five percent) of observed cells. Exact growth conditions that promote sporulation, if any, are currently unknown for *S. arabinophila* DSMZ 106076^T^, but would further help to confirm the capability of this strain to form spores.

### Differences between *S. arabinophila* TLL-A1 and ASF 502

Comparative genome analyses of *S. arabinophila* strains TLL-A1 and ASF 502 revealed a number of distinct differences between the two microbial strains. Some of these predicted functional differences are apparent from the CAZyme repertoire. The genome of strain TLL-A1 harbors a substantially higher number of these CAZyme-encoding genes across all four different categories (CBMs, CEs, GHs, and GTs) than the stain ASF 502. However, the biggest numerical difference between the two strains exists in glycosyl hydrolases. While both strains share most of the detected glycosyl hydrolases (Figure 4), the enrichment for some of the GH in either strain may represent a specific adaptation to an ecological niche. Overall, these differences and those for other predicted molecular functions indicate that the strains may have originated from a common ancestor a long time ago. There is currently limited information on how the genetic repertoire of individual intestinal bacterial strains, such as *Bacteroides thetaiotaomicron* (23), or entire microbial communities change over time. Considering that ASF has been in use for multiple decades and that the comprising strains may have undergone thousands of replications in some mouse colonies, it may seem plausible to assume that substantial genetic changes may have occurred over time. It also needs be noted that strain ASF 502 was isolated from CD-1 mice (derived from Swiss mice) (24), while strain TLL-A1 was isolated from C57BL/6J (derived from C57BL) (25). Both mouse strains (including their progenitors) are genetically distinct, and have been established for over 80 years. This time period may therefore also need to be taken into consideration for a potential evolutionary separation of the two *Schaedlerella* strains as the genetic differences in the microbial strains may represent adaptations to a specific host. Ultimately, it may not be possible to fully tease out the factors that led to genomic differences of two *S. arabinophila* strains, especially considering that the association of their common ancestor with host mice is likely to be older than the history of mice as laboratory model itself. Isolating and characterizing additional *S. arabinophila* strains may help to provide insights into the diversity of this clade, with the resulting pangenome revealing the metabolic potential and flexibility of *S. arabinophila*. In a broader context, understanding strain-specific differences in mouse-associated microorganisms may also increase our understanding of how human-associated microorganisms may adapt to different human individuals and different organs within them.

### Relative abundance of *S. arabinophila* and implications for ASF

Amplicon sequencing was performed on DNA extracted from mouse fecal samples that served as inoculum for the isolation of *S. arabinophila* TLL-A1. The amplicon data reveals that *S. arabinophila* is only present at low relative abundance (less than one percent) in the analysed samples. It needs to be noted that mice were maintained on a standard chow rich in fibre, which generally favours Bacteroidetes over Firmicutes (a phylum *S. arabinophila* belongs to) (26). The latter may be more abundant on other diets with higher caloric intake, such as the western diet (26). It may therefore be misleading to put too much emphasis on the observed relative abundance in this study, as intestinal microbial communities are able to respond quickly and drastically to dietary changes (27). However, it may still be worthwhile to take these observations into consideration when working with ASF.

ASF has served as a valuable synthetic microbial community for a long time, despite some known limitations (28, 29). While ASF 502 may reach high relative abundances in ASF-harbouring mice fed a standard chow (15), it is also apparent from this study that ASF 502 is not a major component of conventionally-raised laboratory mouse microbiota on a polysaccharide rich diet. For future studies it may therefore be necessary to enhance or alter the Altered Schaedler Flora again, taking into consideration also the different experimental conditions, such as dietary changes for the animals. Recent cultivations efforts have resulted in the cultivation of 100 murine intestinal microbes (30). While many more mouse intestinal microorganisms remain to be (re-) cultured, this collection may serve as an important tool to increase the diversity and complexity of autochthonous synthetic microbial communities in mice. In addition, protocols and bioinformatics pipelines, e.g. COPRO-seq (31), have been developed to analyse defined synthetic microbial communities. Combining these tools may help to meet the growing need to develop synthetic communities of autochthonous microorganism that realistically mirror the composition of the conventional mouse microbiota and that allow to study the specific interactions between hosts and microorganisms (32, 33).

In conclusion, this study provides a detailed description of *Schaedlerella arabinophila*, a representative of a new bacterial genus from the intestinal tract of mice and a close relative of *Clostridium* species ASF 502. The results highlight the need to further investigate intraspecies differences of intestinal bacteria and how host-strain specific adaptations may have shaped the genomes of gut commensals.

### Description of *Schaedlerella* gen. nov

Schaedlerella (Schaed.ler.el.la; N.L. fem. n. named after Russel W. Schaedler, one of the pioneering researchers in the study of host-microbiota interactions). The genus represent two strains of a species that are closely related to *Ruminococcus* species and *Clostridium glycyrrhizinilyticum*. Cells are uniform, rod-shaped (2-5 μm long), Gram positive and obligate anaerobe. Saccharoclastic chemo-organotrophs. Peptidoglycan is A1y meso-Dpm-direct type. GC-content inferred from the genomes of strains TLL-A1 and ASF 502 is 47.9%. The type species is *Schaedlerella arabinophila*.

Description of *S. arabinophila* sp. nov. *arabinophila* (a.ra.bi.no.phi.la), N.L. fem. –arabin - natural rubber that has high content of arabinose, adj. philus -liking). Grows on sucrose, D-cellubiose, D-maltose, D-melibiose, D-raffinose, A-lactose, D-mannitol, Glycogen (type II), tannic acids, dextrin, maltodextrin, D-fructose, D-glucose, D-xylose, D-galactose, L-fucose, D-ribose, meso-erythritol, N-acetyl-D-glucosamine, N-acetyl-neuramine, D-arabinose, L-arabinose. No growth was observed (within 15 days) on ethanolamine, choline, starch, cellulose, chitosan, pyruvate. The type strain is DSMZ 106076^T^ / KCTCC 15657^T^.

## Materials and Methods

### Isolation of *S. arabinophila*

*S. arabinophila* TLL-A1 was isolated from freshly collected feces of C57BL/6J mice. Experimental procedures involving animals were performed according to the approved IACUC protocol TLL-16–016. Mouse fecal material was homogenized and serially diluted prior to plating. The medium composition for isolation and maintenance of *S. arabinophila* are as follows (weight given in grams per litre of distilled water): Eight grams sodium acetate (Sigma-Aldrich, St. Louis, Missouri, USA), 0.2g magnesium sulfate heptahydrate (Sigma-Aldrich, Bangalore, India), 0.25g ammonium sulfate (Sigma-Aldrich, Tokyo, Japan), 0.69g potassium phosphate dibasic (Sigma-Aldrich, St. Louis, Missouri, USA), 0.5g yeast extract (Oxoid Ltd., Basingstoke, United Kingdom), 0.5ml of 0.2% resazurin (Sigma, St. Louis, Missouri, USA), 7.5g D-arabinose (Alfa Aesar, Lancashire, United Kingdom), ten ml of trace mineral supplement (ATCC, Manassas, Virginia, USA), and one ml of vitamin supplement (ATCC, Manassas, Virginia, USA). The pH is adjusted to pH7.0 using 37% fuming hydrochloric acid (Merck, Darmstadt, Germany). The prepared medium was filter sterilised and placed into the anaerobic chamber. Filter-sterilized 50 mg of sodium dithionite (Sigma-Aldrich, Munich, Germany) and three grams of sodium bicarbonate (Sigma-Aldrich, St. Louis, Missouri, USA) were further added to the medium at least one day prior to usage. Solid medium, used for streaking and picking out individual colonies to ensure pure culture, was prepared by combining autoclaved 4% agar in distilled water with 500 ml of two-fold concentrated filtered liquid medium. *S. arabinophila* was maintained in the above mentioned liquid medium. 0.5ml of bacteria culture is inoculated into ten ml of fresh medium every two days. Cultures were kept in an anaerobic Balch tube in 37°C incubator. For long-term preservation cultures were stored in 20% glycerol at −80°C.

### Growth characteristics of *S. arabinophila*

Arabinose was omitted from the aforementioned cultivation medium to investigate growth of *S. arabinophila* on different carbon sources. The following carbon sources were tested: sucrose (Nacalai Tesque, Kyoto, Japan), D-cellubiose (Sigma, Dorset, England, United Kingdom), D-maltose monohydrate (Sigma-Aldrich, St. Louis, Missouri, USA), ethanolamine (Sigma-Aldrich, St. Louis, Missouri, USA), choline chloride (Sigma-Aldrich, Shanghai, China), D-melibiose (Sigma-Aldrich, Bratislava, Slovakia), D-raffinose pentahydrate (Sigma-Aldrich, Shanghai, China), L-Lactic acid sodium (Sigma, Buchs, Switzerland), A-lactose monohydrate (Sigma, Munich, Germany), D-mannitol (Sigma-Aldrich, Shanghai, China), glycogen Type II (Sigma, Tokyo, Japan), soluble starch (Sigma-Aldrich, Munich, Germany), tannic acid (Sigma-Aldrich, Shanghai, China), A-cellulose bioreagent (Sigma, St. Louis, Missouri, USA), chitosan (Aldrich, Reykjavik, Iceland), dextrin 10 (Sigma, Shanghai, China), maltodextrin (Aldrich, St. Louis, Missouri, USA), D-fructose (Sigma, St. Louis, Missouri, USA), D-glucose (Sigma, Lyon, France), D-Xylose (Sigma-Aldrich, Shanghai, China), D-galactose (Sigma-Aldrich, Munich, Germany), L-fucose (Sigma, Bratislava, Slovakia), D-ribose (Sigma, Shanghai, China), pyruvic acid sodium (Sigma, Tokyo, Japan), meso-erythritol (Sigma, Shanghai, China), N-acetyl-D-glucosamine (Sigma, Shanghai, China), and N-acetylneuraminic acid (Sigma-Aldrich, St. Louis, Missouri, USA). Each carbon source was added at a final concentration of 10mM except for polymers that were added at 5g/L

To prepare the *S. arabinophila* for inoculation into the various carbon source medium, two sets of four ml of overnight *S. arabinophila* culture was each inoculated into 80 ml of base medium in sterile glass bottles. Cultures were kept in a 37°C incubator overnight before inoculation into the various carbon source medium. Three replicates of each type of medium were made. One ml of two days post-inoculation culture of *S. arabinophila* growing in base medium was inoculated into each of the 20 ml of various carbon source medium. Optical density was visually classified every 24 hours for fifteen days. On the sixth and fifteenth day post inoculation, Optical density was measured using Ultrospec 8000 5706 V1.0.1, at wavelength 600.0nm, bandwidth of 1nm, integration time of one second. For the cultures with sufficient growth, gDNA of the resultant cultures were extracted. The gDNA was then sequenced to ensure that the growth observed was that of *S. arabinophila*.

### Gram and spore stains

Gram reactivity of *S. arabinophila* was investigated using Gram’s crystal violet solution, Gram’s iodine solution, Gram’s safranin solution (Sigma-Aldrich, St. Louis, Missouri, USA) and a decoloriser solution of 1:1 ethanol (Fisher Scientific U.K. Limited, Leicester, United Kingdom) to acetone (Merck Specialities Private Limited, Mumbai, India). Protocol described by Sigma-Aldrich Gram Staining Kit was used. Spore stain was conducted on *S. arabinophila* using Schaeffer & Fulton’s Spore Stain A (Malachite green 50 g/l in water) and Schaeffer & Fulton’s Spore Stain B (Safranin O 5 g/l in water) (Sigma-Aldrich, Buchs, Switzerland). Spore stain protocol used was as described by Sigma-Aldrich Schaeffer and Fulton Spore Stain Kit. The stained cells were imaged using a widefield upright microscope. The camera used was Leica microscope camera DFC7000T and cooled colour camera for LASX. The microscope used is the Axioplan 2 imaging upright motorised microscope. Leica application suite X 3.0.1.15878 from Leica microsystems was used to capture and process the images.

### Characterization of cellular properties

The following characterizations of *S. arabinophila* were performed at DSMZ, Braunschweig, Germany: Analysis of polar lipids, analysis of respiratory quinones, analysis of the cellular fatty acids, peptidoglycan structure, BioMérieux API 50 CH (bioMérieux, Craponne France), BioMérieux API Rapid ID 32 A (bioMérieux, Craponne France), and identification of whole cell sugars.

### DNA extraction and sequencing

Liquid culture of *S. arabinophila* was grown to optical density of 1.80 at a wavelength of 600nm. 50 ml culture was centrifuged (Beckman Coulter rotor JA10, 6000g, 20min, 4°C) to pellet the cells out of the media and subject to genomic DNA extraction (modified after (34)) using two rounds of phenol-chloroform purification. The extracted gDNA was stored in −20 °C until further use. Ten μg of prepared genomic DNA was purified with AMPure XP magnetic beads and quality checked with Nano drop and Qubit fluorometer (Invitrogen, Carlsbad, CA, USA) and subsequently sent for PacBio RSII Single Molecule and Illumina MiSeq paired-end (251×251bp) sequencing at Singapore Centre for Environmental Life Sciences Engineering (SCELSE) in Nanyang Technological University.

### Genome assembly and annotation

Raw PacBio sequencing reads were *de novo* assembled with SMRT analysis software (via HGAP 4) with standard parameters except the estimated genome size was set as 6.5 Mb according to the genome of ASF 502 (12). Reads from MiSeq paired-end sequencing were used for further error correction using Pilon (17).

As an initial quality check step of the assembly, RNAmmer (35) was used to annotate and verify the RNA genes, of which the 16S rRNA gene was further used for phylogenetic tree construction. In addition, CheckM (36) was used to assess genome completeness, percentage contamination, as well as finding out missing single-copy marker genes. Because the polished genome consisted of three contigs of which two were short (6.35 Mb, 0.16 Mb and 9.5 kb), Plasmid Finder (16) was used to assess if any were plasmids. Genomes were annotated using RASTtk (37) online through PATRIC server (38).

### Phylogeny

16S rRNA gene and whole genome phylogenies were constructed in order to compare the evolutionary placement of strain TLL-A1. The 16S rRNA gene was extracted from the polished genome using RNAmmer as described earlier, aligned with SINA (39) and phylogeny constructed by RAxML within ARB (40). Whole genome phylogeny was constructed in PATRIC using conserved protein sequences via RAxML. Two genomes were further compared by average amino acid identity (AAI) (41) and *in silico* DNA-DNA hybridization (by genome sequence-based delineation (GGDC)) (42) to determine if the two strains belong to the same species. Typically, genomes are considered to belong to the same species for AAAI values above 95% or over 70% DNA-DNA hybridization.

### Metabolic reconstruction and detection of potential pathogenic activity

Further downstream analysis, including genome visualization, comparison and metabolic construction, was conducted in PATRIC (38). The genome visualization was conducted using circular display where genome contiguity, CDS, non-CDS features, antimicrobial resistance (AMR) genes, transporters, GC content and GC skew were displayed. Further metabolic potential of TLL-A1 was determined by PATRIC interface, specifically looking into presence/absence of key enzymes in some instances. In brief, major central metabolism genes (i.e. Embden-Meyerhof-Parnas pathway, citric acid cycle, pentose phosphate pathway, NADH-dehydrogenase, cytochrome C, cytochrome C oxidase, ATP synthase genes) were checked. Potential genomic islands (GI) were visualized using IslandViewer4 (43) and gene annotation for major GI regions were checked.

### Genome Comparison

The assembly was compared for similarities and differences with the previously sequenced draft genome *Clostridium* sp. ASF 502. OrthoVenn (44) was used for comparisons and annotation of orthologous gene clusters. Given the similarity of the two genomes, COGs that were unique to each genome were further compared according to gene ontology (GO) classification. Additionally, carbohydrate-active enzymes (CAZymes), important for carbohydrate breakdown, were compared between the two genomes. CAZymes were annotated via dbCAN2 meta server (45) and the major class of CAZy checked against the online database (46). Note that only CAZymes identified with > 2 methods (out of HMMER (E-Value < 1e-15, coverage > 0.35; http://hmmer.org/), DIAMOND (E-Value < 1e-102; (47)) and Hotpep (Frequency > 2.6, Hits > 6; (48)) were considered, as recommended by the authors.

### DNA extraction and library preparation for amplicon sequencing

Samples were transferred to sterile bead-beating vials containing ~0.7g zirconium beads and 200μl of 20% SDS. 282μl Buffer A (see Rius *et al* for composition (49)), 268μl Buffer PM, 550μl phenol/chloroform/isoamyl alcohol (25:24:1, pH 8) were added. The pelleted cells were subject to phenol-chloroform-based extraction, after combined mechanical and chemical lysis of cells (49). The samples were quantified and diluted to 40 ng/μl per sample and stored in −20 °C for further use.

The extracted DNA was processed according to the Earth Microbiome Project (http://www.earthmicrobiome.org/protocols-and-standards/16s/) as described in Thompson *et al* [13] except a dual-indexing strategy was used as described by (50) to reduce the number of barcoded primers required. PCR was conducted with QIAGEN Taq MasterMix (CAT NO 201445, QIAGEN, Germany) in triplicates (plus one negative control per sample) under the condition: 94 °C for 3 mins; 35 cycles of 94 °C for 45 secs, 50 °C for 60 °C, 72 °C for 90 secs; 72 °C for 10 mins; 4 °C hold. The amplicons were quantified with Quant IT Picogreen (CAT NO P7589, Thermo Fisher Scientific, USA), pooled together in equimolar concentrations and sequenced at Singapore Centre for Environmental Life Sciences Engineering (SCELSE) at Nanyang Technological University using Illumina MiSeq paired-end chemistry (251×251bp). The output raw sequences were analysed with MOTHUR v1.39.1 (51) according to the standard MiSeq protocol (52). The sequence pairs were merged, de-multiplexed and quality-filtered accordingly. Furthermore, because Lachnospiraceae UCG-006 had the highest sequence identity to *S. arabinophila* at 16S rRNA level, all sequences that were classified as Lachnospiraceae UCG-006 were picked out and reclassified with a modified database that contained the *S. arabinophila* 16S rRNA gene (annotated by RNAmmer from the genome sequenced in this study) to determine the relative abundance of the latter. Visualization of processed data was conducted by R project following tutorials provided in https://joey711.github.io/phyloseq/index.html. Packages used include: ape (53), dplyr (54), ggdendro (55), ggplot2 (56), ggpubr (57), gplots (58), grid (59), gridExtra (60), Heatplus (61), pheatmap (6), phyloseq (62), plyr (63), RColorBrewer (64), reshape2 (65), sfsmisc (66), vegan (67) and viridis (68).

### Accession numbers

The demultiplexed, pair-matched amplicon sequences are deposited in NCBI SRA under accession number PRJNA498133. The assembled genome of *Schaedlerella arabinophila* TLL-A1 is deposited in GenBank under the accession number PRJNA497957.

## Acknowledgments

We thank Daniela Moses at SCELSE Singapore for performing the PacBio sequencing, and Amit Anand and Muhammad Khairillah Bin Nanwi at TLL biocomputing for bioinformatics support. This work was funded by Temasek Life Sciences Laboratory.

## References

1. Shreiner AB, Kao JY, Young VB. 2015. The gut microbiome in health and in disease. Current opinion in gastroenterology 31:69.

2. Wade WG. 2013. The oral microbiome in health and disease. Pharmacological research 69:137–143.

3. Eckburg PB, Bik EM, Bernstein CN, Purdom E, Dethlefsen L, Sargent M, Gill SR, Nelson KE, Relman DA. 2005. Diversity of the human intestinal microbial flora. science 308:1635–1638.

4. Lukeš J, Stensvold CR, Jirků-Pomajbíková K, Parfrey LW. 2015. Are human intestinal eukaryotes beneficial or commensals? PLoS pathogens 11:e1005039.

5. Gordon HA, Pesti L. 1971. The gnotobiotic animal as a tool in the study of host microbial relationships. Bacteriological reviews 35:390.

6. Biggs MB, Medlock GL, Moutinho TJ, Lees HJ, Swann JR, Kolling GL, Papin JA. 2017. Systems-level metabolism of the altered Schaedler flora, a complete gut microbiota. The ISME journal 11:426.

7. Bouskra D, Brézillon C, Bérard M, Werts C, Varona R, Boneca IG, Eberl G. 2008. Lymphoid tissue genesis induced by commensals through NOD1 regulates intestinal homeostasis. Nature 456:507.

8. Smith PM, Howitt MR, Panikov N, Michaud M, Gallini CA, Bohlooly-y M, Glickman JN, Garrett WS. 2013. The microbial metabolites, short-chain fatty acids, regulate colonic Treg cell homeostasis. Science 341:569–573.

9. Orcutt R, Gianni F, Judge R. 1987. Development of an “Altered Schaedler Flora” for NCI gnotobiotic rodents. Microecol Ther 17.

10. Schaedler RW, Dubos R, Costello R. 1965. The development of the bacterial flora in the gastrointestinal tract of mice. Journal of Experimental Medicine 122:59–66.

11. Dewhirst FE, Chien C-C, Paster BJ, Ericson RL, Orcutt RP, Schauer DB, Fox JG. 1999. Phylogeny of the defined murine microbiota: altered Schaedler flora. Applied and environmental microbiology 65:3287–3292.

12. Wannemuehler MJ, Overstreet A-M, Ward DV, Phillips GJ. 2014. Draft genome sequences of the altered schaedler flora, a defined bacterial community from gnotobiotic mice. Genome announcements 2:e00287–00214.

13. Robertson BR, O’Rourke JL, Neilan BA, Vandamme P, On SL, Fox JG, Lee A. 2005. Mucispirillum schaedleri gen. nov., sp. nov., a spiral-shaped bacterium colonizing the mucus layer of the gastrointestinal tract of laboratory rodents. International journal of systematic and evolutionary microbiology 55:1199–1204.

14. Loy A, Pfann C, Steinberger M, Hanson B, Herp S, Brugiroux S, Neto JCG, Boekschoten MV, Schwab C, Urich T. 2017. Lifestyle and horizontal gene transfer-mediated evolution of Mucispirillum schaedleri, a core member of the murine gut microbiota. Msystems 2:e00171–00116.

15. Sarma-Rupavtarm RB, Ge Z, Schauer DB, Fox JG, Polz MF. 2004. Spatial distribution and stability of the eight microbial species of the altered schaedler flora in the mouse gastrointestinal tract. Applied and environmental microbiology 70:2791–2800.

16. Carattoli A, Zankari E, Garcia-Fernandez A, Voldby Larsen M, Lund O, Villa L, Moller Aarestrup F, Hasman H. 2014. In silico detection and typing of plasmids using PlasmidFinder and plasmid multilocus sequence typing. Antimicrob Agents Chemother 58:3895–3903.

17. Walker BJ, Abeel T, Shea T, Priest M, Abouelliel A, Sakthikumar S, Cuomo CA, Zeng Q, Wortman J, Young SK, Earl AM. 2014. Pilon: an integrated tool for comprehensive microbial variant detection and genome assembly improvement. PLoS One 9:e112963.

18. El Kaoutari A, Armougom F, Gordon JI, Raoult D, Henrissat B. 2013. The abundance and variety of carbohydrate-active enzymes in the human gut microbiota. Nature reviews Microbiology 11:497.

19. Flint HJ, Scott KP, Duncan SH, Louis P, Forano E. 2012. Microbial degradation of complex carbohydrates in the gut. Gut microbes 3:289–306.

20. Martens EC, Koropatkin NM, Smith TJ, Gordon JI. 2009. Complex glycan catabolism by the human gut microbiota: the Bacteroidetes Sus-like paradigm. Journal of Biological Chemistry:jbc. R109. 022848.

21. LeBlanc DJ, Mortlock RP. 1971. Metabolism of D-arabinose: a new pathway in Escherichia coli. Journal of bacteriology 106:90–96.

22. Becker DJ, Lowe JB. 2003. Fucose: biosynthesis and biological function in mammals. Glycobiology 13:41R–53R.

23. Peterson DA, Planer JD, Guruge JL, Xue L, Downey-Virgin W, Goodman AL, Seedorf H, Gordon JI. 2015. Characterizing the interactions between a naturally-primed immunoglobulin A and its conserved Bacteroides thetaiotaomicron species-specific epitope in gnotobiotic mice. Journal of Biological Chemistry:jbc. M114. 633800.

24. Chia R, Achilli F, Festing MF, Fisher EM. 2005. The origins and uses of mouse outbred stocks. Nature genetics 37:1181.

25. Morse HC. 1978. Origins of inbred mice. Academic Press.

26. Jumpertz R, Le DS, Turnbaugh PJ, Trinidad C, Bogardus C, Gordon JI, Krakoff J. 2011. Energy-balance studies reveal associations between gut microbes, caloric load, and nutrient absorption in humans–. The American journal of clinical nutrition 94:58–65.

27. David LA, Maurice CF, Carmody RN, Gootenberg DB, Button JE, Wolfe BE, Ling AV, Devlin AS, Varma Y, Fischbach MA, Biddinger SB, Dutton RJ, Turnbaugh PJ. 2014. Diet rapidly and reproducibly alters the human gut microbiome. Nature 505:559.

28. Norin E, Midtvedt T. 2010. Intestinal microflora functions in laboratory mice claimed to harbor a “normal” intestinal microflora. Is the SPF concept running out of date? Anaerobe 16:311–313.

29. Wymore Brand M, Wannemuehler MJ, Phillips GJ, Proctor A, Overstreet A-M, Jergens AE, Orcutt RP, Fox JG. 2015. The altered Schaedler flora: continued applications of a defined murine microbial community. ILAR journal 56:169–178.

30. Lagkouvardos I, Pukall R, Abt B, Foesel BU, Meier-Kolthoff JP, Kumar N, Bresciani A, Martínez I, Just S, Ziegler C, Brugiroux S, Garzetti D, Wenning M, Bui TPN, Wang J, Hugenholtz F, Plugge CM, Peterson DA, Hornef MW, Baines JF, Smidt H, Walter J, Kristiansen K, Nielsen HB, Haller D, Overmann J, Stecher B, Clavel T. 2016. The Mouse Intestinal Bacterial Collection (miBC) provides host-specific insight into cultured diversity and functional potential of the gut microbiota. Nature microbiology 1:16131.

31. McNulty NP, Yatsunenko T, Hsiao A, Faith JJ, Muegge BD, Goodman AL, Henrissat B, Oozeer R, Cools-Portier S, Gobert G, Chervaux C, Knights D, Lozupone CA, Knight R, Duncan AE, Bain JR, Muehlbauer MJ, Newgard CB, Heath AC, Gordon JI. 2011. The impact of a consortium of fermented milk strains on the gut microbiome of gnotobiotic mice and monozygotic twins. Science translational medicine 3:106ra106–106ra106.

32. Chung H, Pamp SJ, Hill JA, Surana NK, Edelman SM, Troy EB, Reading NC, Villablanca EJ, Wang S, Mora JR, Umesaki Y, Mathis D, Benoist C, Relman DA, Kasper DL. 2012. Gut immune maturation depends on colonization with a host-specific microbiota. Cell 149:1578–1593.

33. Seedorf H, Griffin NW, Ridaura VK, Reyes A, Cheng J, Rey FE, Smith MI, Simon GM, Scheffrahn RH, Woebken D. 2014. Bacteria from diverse habitats colonize and compete in the mouse gut. Cell 159:253–266.

34. Sambrook J, Fritsch EF, Maniatis T. 1989. Molecular cloning: a laboratory manual. Cold spring harbor laboratory press.

35. Lagesen K, Hallin P, Rødland EA, Stærfeldt H-H, Rognes T, Ussery DW. 2007. RNAmmer: consistent and rapid annotation of ribosomal RNA genes. Nucleic Acids Research 35:3100–3108.

36. Parks DH, Imelfort M, Skennerton CT, Hugenholtz P, Tyson GW. 2015. CheckM: assessing the quality of microbial genomes recovered from isolates, single cells, and metagenomes. Genome research:gr.186072.186114.

37. Brettin T, Davis JJ, Disz T, Edwards RA, Gerdes S, Olsen GJ, Olson R, Overbeek R, Parrello B, Pusch GD, Shukla M, Thomason III JA, Stevens R, Vonstein V, Wattam AR, Xia F. 2015. RASTtk: a modular and extensible implementation of the RAST algorithm for building custom annotation pipelines and annotating batches of genomes. Scientific reports 5:8365.

38. Wattam AR, Davis JJ, Assaf R, Boisvert S, Brettin T, Bun C, Conrad N, Dietrich EM, Disz T, Gabbard JL, Gerdes S, Henry CS, Kenyon RW, Machi D, Mao C, Nordberg EK, Olsen GJ, Murphy-Olson DE, Olson R, Overbeek R, Parrello B, Pusch GD, Shukla M, Vonstein V, Warren A, Xia F, Yoo H, Stevens RL. 2017. Improvements to PATRIC, the all-bacterial Bioinformatics Database and Analysis Resource Center. Nucleic Acids Research 45:D535–D542.

39. Pruesse E, Peplies J, Glöckner FO. 2012. SINA: accurate high-throughput multiple sequence alignment of ribosomal RNA genes. Bioinformatics 28:1823–1829.

40. Ludwig W, Strunk O, Westram R, Richter L, Meier H, Yadhukumar, Buchner A, Lai T, Steppi S, Jobb G, Foerster W, Brettske I, Gerber S, Ginhart AW, Gross O, Grumann S, Hermann S, Jost R, König A, Liss T, Lüßmann R, May M, Nonhoff B, Reichel B, Strehlow R, Stamatakis A, Stuckmann N, Vilbig A, Lenke M, Ludwig T, Bode A, Schleifer KH. 2004. ARB: a software environment for sequence data. Nucleic acids research 32:1363–1371.

41. Goris J, Konstantinidis KT, Klappenbach JA, Coenye T, Vandamme P, Tiedje JM. 2007. DNA–DNA hybridization values and their relationship to whole-genome sequence similarities. International journal of systematic and evolutionary microbiology 57:81–91.

42. Meier-Kolthoff JP, Auch AF, Klenk H-P, Göker M. 2013. Genome sequence-based species delimitation with confidence intervals and improved distance functions. BMC bioinformatics 14:60.

43. Bertelli C, Laird MR, Williams KP, Group SFURC, Lau BY, Hoad G, Winsor GL, Brinkman FS. 2017. IslandViewer 4: expanded prediction of genomic islands for larger-scale datasets. Nucleic acids research 45:W30–W35.

44. Wang Y, Coleman-Derr D, Chen G, Gu YQ. 2015. OrthoVenn: a web server for genome wide comparison and annotation of orthologous clusters across multiple species. Nucleic Acids Research 43:W78–W84.

45. Zhang H, Yohe T, Huang L, Entwistle S, Wu P, Yang Z, Busk PK, Xu Y, Yin Y. 2018. dbCAN2: a meta server for automated carbohydrate-active enzyme annotation. Nucleic Acids Research 46:W95–W101.

46. Lombard V, Golaconda Ramulu H, Drula E, Coutinho PM, Henrissat B. 2014. The carbohydrate-active enzymes database (CAZy) in 2013. Nucleic Acids Research 42:D490–D495.

47. Buchfink B, Xie C, Huson DH. 2014. Fast and sensitive protein alignment using DIAMOND. Nature methods 12:59.

48. Busk PK, Pilgaard B, Lezyk MJ, Meyer AS, Lange L. 2017. Homology to peptide pattern for annotation of carbohydrate-active enzymes and prediction of function. BMC bioinformatics 18:214.

49. Rius A, Kittelmann S, Macdonald K, Waghorn G, Janssen P, Sikkema E. 2012. Nitrogen metabolism and rumen microbial enumeration in lactating cows with divergent residual feed intake fed high-digestibility pasture. Journal of dairy science 95:5024–5034.

50. Fadrosh DW, Ma B, Gajer P, Sengamalay N, Ott S, Brotman RM, Ravel J. 2014. An improved dual-indexing approach for multiplexed 16S rRNA gene sequencing on the Illumina MiSeq platform. Microbiome 2:6.

51. Schloss PD, Westcott SL, Ryabin T, Hall JR, Hartmann M, Hollister EB, Lesniewski RA, Oakley BB, Parks DH, Robinson CJ, Sahl JW, Stres B, Thallinger GG, Van Horn DJ, Weber CF. 2009. Introducing mothur: open-source, platform-independent, community-supported software for describing and comparing microbial communities. Applied and environmental microbiology 75:7537–7541.

52. Kozich JJ, Westcott SL, Baxter NT, Highlander SK, Schloss PD. 2013. Development of a dual-index sequencing strategy and curation pipeline for analyzing amplicon sequence data on the MiSeq Illumina sequencing platform. Applied and environmental microbiology:AEM. 01043–01013.

53. Paradis E, Schliep K, Schwartz R. 2018. ape 5.0: an environment for modern phylogenetics and evolutionary analyses in R. Bioinformatics 1:3.

54. Wickham H, Francois R, Henry L, Müller K. 2015. dplyr: A grammar of data manipulation. R package version 04 3.

55. de Vries A, Ripley B. 2013. Ggdendro: tools for extracting dendrogram and tree diagram plot data for use with ggplot. R package version 01-12, Retrieved from http://CRANR-projectorg/package=ggdendro.

56. Wickham H. 2016. ggplot2: elegant graphics for data analysis. Springer.

57. Kassambara A. 2017. ggpubr:“ggplot2” based publication ready plots. R package version 01 6.

58. Warnes GR, Bolker B, Bonebakker L, Gentleman R, Huber W, Liaw A, Lumley T, Maechler M, Magnusson A, Moeller S. 2009. gplots: Various R programming tools for plotting data. R package version 2:1.

59. Murrell P. 2016. R graphics. CRC Press.

60. Auguie B. 2012. gridExtra: functions in Grid graphics. R package version 09 1.

61. Ploner A. 2012. Heatplus: Heatmaps with row and/or column covariates and colored clusters. R package version 2.

62. McMurdie PJ, Holmes S. 2013. phyloseq: an R package for reproducible interactive analysis and graphics of microbiome census data. PloS one 8:e61217.

63. Wickham H. 2009. plyr: Tools for splitting, applying and combining data. R package version 01 9:651.

64. Neuwirth E. 2011. Package ‘RColorBrewer’. CRAN 2011-06-17 08: 34: 00. Apache License 2.0,

65. Wickham H. 2012. reshape2: Flexibly Reshape Data: A Reboot of the Reshape Package. R Package Version 1.

66. Maechler M. 2012. sfsmisc: Utilities from seminar fuer Statistik ETH Zurich. R package version:1.0-20.

67. Oksanen J, Blanchet FG, Kindt R, Legendre P, O’hara R, Simpson GL, Solymos P, Stevens MHH, Wagner H. 2010. vegan: Community Ecology Package. R package version 1.17-2. http://cranr-projectorg> Acesso em 23:2010.

68. Jovel J, Patterson J, Wang W, Hotte N, O’Keefe S, Mitchel T, Perry T, Kao D, Mason AL, Madsen KL, Wong GK-S. 2016. Characterization of the gut microbiome using 16S or shotgun metagenomics. Frontiers in microbiology 7:459.

